# Scale free topology as an effective feedback system

**DOI:** 10.1101/696575

**Authors:** Alexander Rivkind, Hallel Schreier, Naama Brenner, Omri Barak

## Abstract

Biological networks are often heterogeneous in their connectivity pattern, with degree distributions featuring a heavy tail of highly connected hubs. The implications of this heterogeneity on dynamical properties are a topic of much interest. Here we introduce a novel approach to analyze such networks the lumped hub approximation. Based on the observation that in finite networks a small number of hubs have a disproportionate effect on the entire system, we construct an approximation by lumping these nodes into a single effective hub, and replacing the rest by a homogeneous bulk. We use this approximation to study dynamics of networks with scale-free degree distributions, focusing on their probability of convergence to fixed points. We find that the approximation preserves convergence statistics over a wide range of settings. Our mapping provides a parametrization of scale free topology which is predictive at the ensemble level and also retains properties of individual realizations. Specifically for *outgoing* scale-free distributions, the role of the effective hub on the network can be elucidated by feedback analysis. We show that outgoing hubs have an organizing role that can drive the network to convergence, in analogy to suppression of chaos by an external drive. In contrast, incoming hubs have no such property, resulting in a marked difference between the behavior of networks with outgoing vs. incoming scale free degree distribution. Combining feedback analysis with mean field theory predicts a transition between convergent and divergent dynamics which is corroborated by numerical simulations. Our results show how interpreting topology as a feedback circuit can provide novel insights on dynamics. Furthermore, we highlight the effect of a handful of outlying hubs, rather than of the connectivity distribution law as a whole, on network dynamics.

## I. INTRODUCTION

Complex networks, their structure and dynamics, have become an indispensable tool for investigating biological systems [1–3]. Networks provide a mathematical model for a system of many interacting components; such models are a cornerstone of computational neuroscience and are widely used in cell biology to describe genetic networks, protein interactions, metabolism and more. Motivated at least partly by these applications, the mathematical theory of network analysis has gained general and fundamental interest and has advanced tremendously in the past decades. Topics of interest include network structure and topology; dynamic behaviour such as fixed points, their stability and their scaling with network size; robustness of these dynamic properties to noise and to evolution; and network controllability and information processing properties [4–7].

One important property of biological networks that has raised much interest is their heterogeneous topology. Analyses of metabolic, protein interaction, gene regulatory and neural networks all show heterogeneous connectivity distributions, including heavy tails and modular structure [8–14]. Heterogeneous networks, such as those with a broad connectivity distribution, are generally more difficult to analyze; inferring their detailed topology requires exceedingly high statistics. Power-law distributions often provide a good approximation to such networks, but statistical difficulties have led to some debate concerning their adequacy [15].

Understanding how heterogeneous connectivity profiles relate to properties of the associated dynamical system is a non-trivial theoretical challenge [16]. One of the main tools to understand the dynamics of *homogeneous* random networks is the use of mean field approximations [17, 18], which rely on two assumptions – homogeneity and a large network size. Corrections for finite size were made as perturbations around the thermodynamic limit [19]. Heterogeneity was approached by dividing networks into homogeneous sub-networks [20–23]. It might be beneficial to develop a complementary approach that does not rely on the usual mean field approximation or its extensions. We shall see that two common difficulties of realistic, far-from-ideal networks – their heterogeneous structure and finite size – perhaps surprisingly work together to shed light on the relation between their structure and dynamics.

We focus on the probability of random network ensembles to converge to attractors, and how this probability is modulated by ensemble topology. Convergence to fixed points is one of the simplest dynamical behaviors, and hence a suitable starting point for analysis. Moreover, stable fixed points have traditionally been linked to fundamental functional properties of networks [24–26].

We were specifically motivated by a recent observation linking topological properties of gene regulatory networks to their probability of convergence, and to the ability of cells to adapt to challenges [27]. In this work, Schreier and colleagues demonstrated in a random network model, that convergence probability is dramatically larger in networks with Scale-Free Out (SFO) degree distributions than in their transposed counterparts with Scale-Free In (SFI) degrees. This is an intriguing result since the two transposed ensembles of interaction matrices share the same eigenvalue statistics. The difference in the probability of convergent dynamics therefore must depend on properties beyond the spectrum. This is in contrast to homogeneous random network ensembles, where dynamic properties are well predicted from the spectrum (e.g. the transition between chaotic and dissipative dynamics [17]).

Here we develop a novel approximation for network ensembles with a broad connectivity distribution, based on the dominant role of a handful of hubs in a finite network. Identifying such hubs [28], understanding their impact on network functionality [29], and advancing the related theory [30] are all active research areas. By approximating a few hubs at the tail-end of the distribution by a single effective hub, we map the problem of heterogeneous network dynamics to that of a homogeneous network coupled to a single external node: each SF network is approximated by a single *lumped-hub* connected to a homogeneous network of the remaining nodes (the bulk). Despite the simplification, this mapping retains many of the properties of the original network ensemble. The new network is parameterized by hub strength and bulk connectivity.

The lumped-hub ensemble can be analyzed by a combination of mean field approximations and feedback analysis. This analysis predicts a phase transition between converging and diverging dynamics as a function of the balance between hub strength and bulk connectivity, which is verified by numerical simulations. We show that the prediction holds on average for scale-free networks in a range of parameters relevant to real networks. Thus, the role of outgoing hubs can be considered analogous to that of an external input in overcoming recurrent activity, suppressing chaotic dynamics, and ultimately driving the system to a stable fixed point.

The success of our mapping suggests that the existence of dominant hubs and their relative strength is a predictive property of a given network’s dynamics, possibly more so than the precise scaling of the distribution from which the connectivity was sampled. Deviations from the theory stemming from large variability in scale-free ensembles as well as corrections to the mean field assumption are discussed.

Our results offer a novel framework for analyzing heterogeneous networks, which links insights from two different angles – internal dynamics of complex networks and the response to external input of simpler networks. This approach highlights the crucial role of a small number of hubs in a finite network, in contrast to other quantitative features of the connectivity distribution, and identifies robust parameters that affect dynamics.

## II. DYNAMICS WITH SCALE-FREE NETWORKS: MOTIVATING OBSERVATIONS

In what follows we analyze a model of nonlinear dynamics, often used to describe biological interactions such as in neuronal or genetic regulatory networks [24, 31]. Binary dynamic variables, *s*_*i*_ (1 ≤ *i* ≤ *N*), approximate the activity of *N* individual elements as being in one of two states: expressed (+1) or repressed (−1) genes, firing or quiescent neurons. The effect of elements on one another is described by a weighted sum of all incoming links:

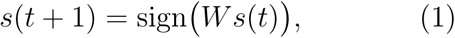

where *W* is the *N* × *N* connectivity matrix with real entries giving the strengths of interactions, drawn from some random network ensemble. It is a special case of the widely studied Boolean networks [24, 32, 33], with the Boolean function chosen here to be the sign of the inputs’ weighted sum.

The ensemble from which *W* is drawn is a crucial ingredient of Eq. (1). To study the effects of topology on the dynamics, we define *W* = *T* ∘ *J* as a Hadamard (element-wise) product between a random topology matrix *T* and a random interaction strength matrix *J*. The 0*/*1 adjacency matrix *T* defines a directed graph, whose edges are sampled from specified distributions of incoming and outgoing connections (see Methods). The strengths *J* are i.i.d. Gaussian variables with zero mean and standard deviation of unity.

For networks described by a directed graph, in general, incoming and outgoing degree distributions, *P*_*in*_(*k*), *P*_*out*_(*k*), need not be the same. In one case of special interest, gene regulatory networks, an empirical observation of such dissimilarity was reported. Outgoing connections were found to have a broad distribution consistent with a power-law: *P*_*out*_(*k*) ∼ *k*^−*γ*^, a scale-free (SF) distribution, while incoming connections were much more narrowly distributed around the average [34]. These statistical properties represent the biological observation that while any given gene is regulated by at most a handful of others, some transcription factors can regulate the expression of up to hundreds of genes in the cell. These ‘master regulators’ are of much interest in cell biology, and have been at the focus of specific molecular biology studies [35].

In a recent work describing gene regulatory networks by random interaction matrices, it was found that networks drawn from ensembles with outgoing hubs have a much larger probability to converge to attractors under exploratory adaptation [27]. Ensembles that showed efficient ability for adaptation also exhibited a high probability of convergence to a fixed point attractor in their intrinsic dynamics.

Particularly noteworthy in this context is the observation that for the transpose random network ensemble, where incoming connections are broadly distributed whereas outgoing connection are not, the abundance of fixed point attractors is decreased dramatically. This result is illustrated for the model of Eq. (1) in Fig. 1A (open circles). For each of the two ensembles – Scale-Free Out degree distribution (SFO) and the transposed SFI – the fraction of simulations converging to a fixed point after a given time interval was computed; this is used as an estimate of the convergence probability across the ensemble. The result is presented as a function of network size for both ensembles. For SFO networks a considerable convergence probability is maintained up to the maximal network size of *N* = 5000, whereas for SFI networks this probability decreases rapidly and is negligible for networks with *N* > 1000. These results are consistent with those of [27], where a model with continuous dynamics were used, extending the observation on the two transposed ensembles to Boolean dynamics.

**FIG. 1.**
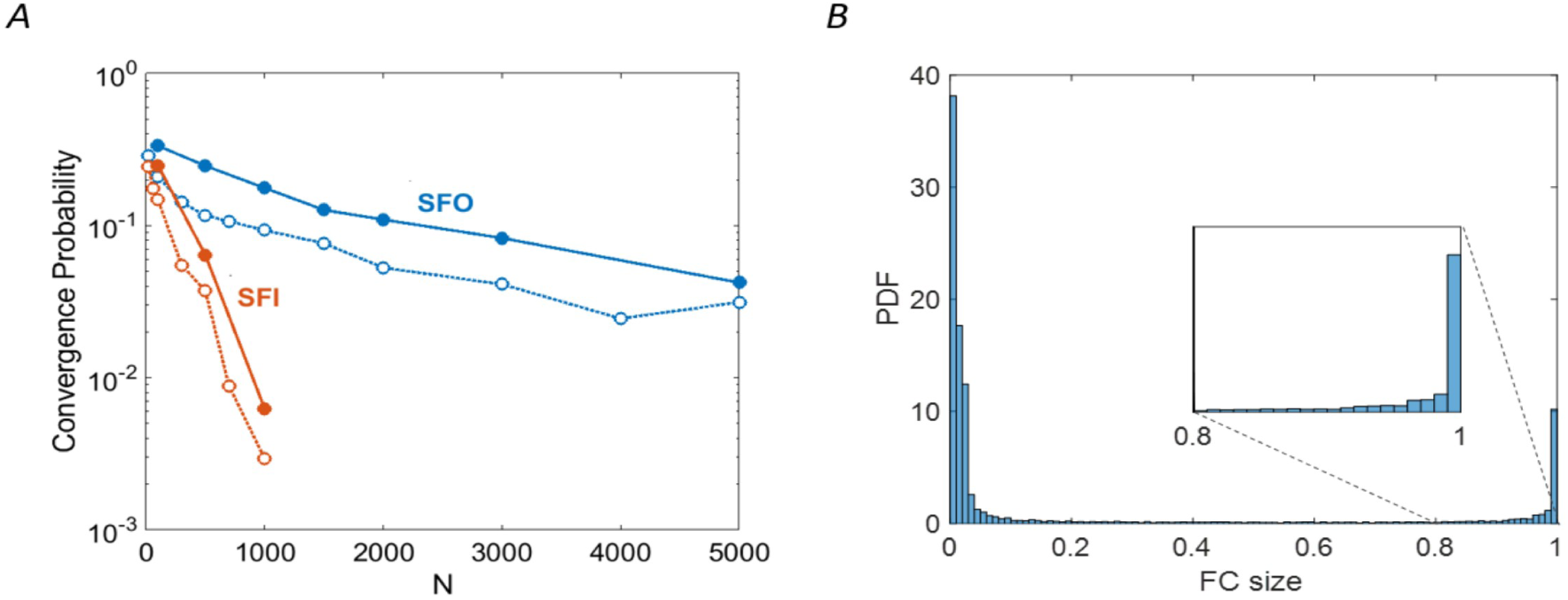
Dynamic properties of Scale-Free-Out (SFO, blue) vs. Scale-Free-In (SFI, red) network ensembles. **(A)** Convergence probability is shown as a function of network size. Filled circles and solid lines show convergence to frozen cores larger than 90% of the network; open circles and dashed lines show convergence to fixed points. Each data point represents an ensemble of 600 networks with random *T, J* and initial conditions. **(B)** Distribution of frozen core sizes (FC) relative to network size, across the SFO ensemble for N=1500. The distribution is further zoomed near FC=1. SF parameters for both panels are *γ* = 2.2 and *k*_*min*_ = 1, Binomial parameters are *N* and *p* = *k/N* with *k* ∼ 5 (exact value of *k* is determined by the realization of SF distribution. See methods for details).

When simulating these Boolean dynamics, we noticed that networks often converge to a state where the vast majority of the nodes are fixed – “frozen”, while the rest continue to change (Fig. 1B). From a biological perspective these are of interest as partially fixed states, and mathematically they have been studied in the context of general Boolean networks [33]. In what follows we define a quasi-fixed-point (QFP) as a state in which most (> 90%) of the nodes are frozen. Fig. 1A (filled circles) demonstrates that the probability of convergence to a QFP behaves qualitatively similar to the corresponding probability for fixed points over the two ensembles: the remarkable difference between SFO and SFI ensembles and its dependence on network size remain the same. This shows that the constraining effect of outgoing hubs on network dynamics is not a unique feature of fixed points, but is also present for weaker notions of convergence.

Our empirical observations rely on convergence of dynamics – and hence is affected both by the existence of fixed points and by their stability. In Appendix C we show that networks from both SFO and SFI ensembles typically have *O*(1) fixed points. This suggests that the reported differences between ensembles stem from stability of fixed points. In the following sections we develop an approach which enables to better understand the stability properties of the two ensembles.

## III. THE LUMPED-HUB APPROXIMATION

A finite realization of a power-law connectivity distribution is dominated by a handful of hubs connected to a macroscopic fraction of the network. Fig. 2A shows an empirical histogram of the outgoing connections in an SFO network with *N* = 1500, where the largest hubs each connect to ∼ 1000 nodes. To capture the properties of such a network we divide it into a small group of leading hubs, and the “bulk” – the rest of the network. The *m* largest hubs are lumped into one node preserving its connection to the bulk; the remaining nodes are substituted by a homogeneous network with binomially distributed incoming and outgoing degrees, preserving the average connectivity. We call this mapping the “lumped hub approximation”; the resulting network is characterized by the bulk mean connectivity *k*_*b*_ and the pattern of connections from the hub, which depend on both the original network topology and the lumping parameter *m*.

**FIG. 2.**
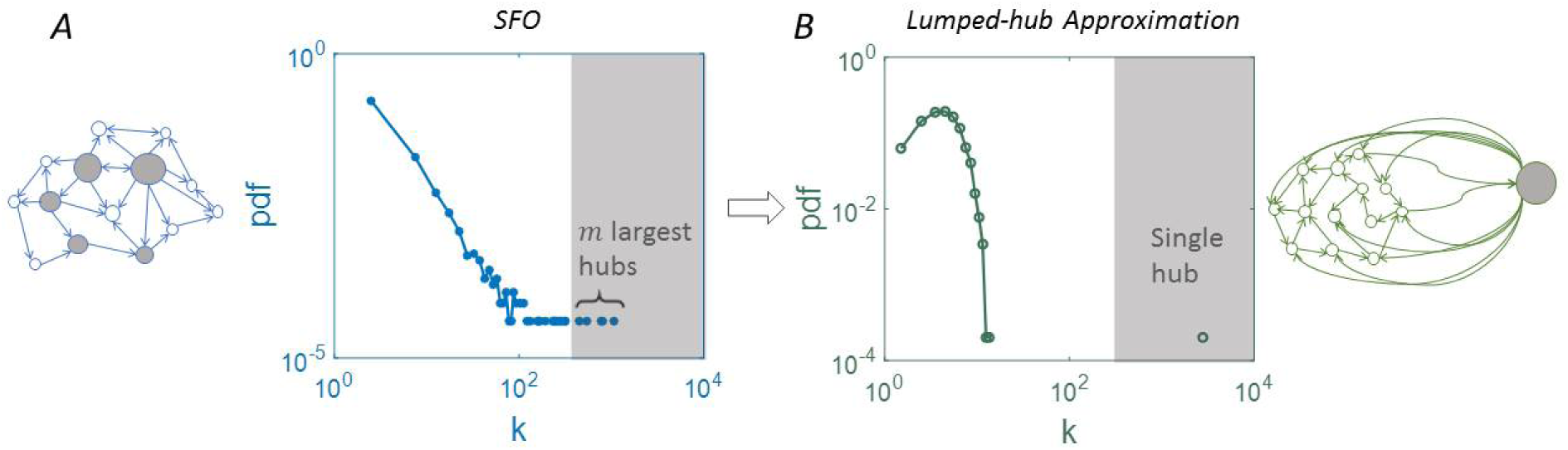
The lumped-hub approximation. (A) In a finite network with SFO degree distribution, the *m* largest hubs (gray shading) are connected to a macroscopic fraction of the network. The histogram shows the number of outgoing connections from each node. (B) In the lumped-hub approximation of the same network, these are substituted by one effective hub and the rest of the nodes are approximated by a Binomial network. The connection strengths of the single hub to the bulk are determined by summing the strengths of the *m* lumped hubs. The mean degree of the bulk, *k*_*b*_, is retained in the mapping.

Despite the crudeness of the lumping approximation, we find that it preserves the dynamic properties of SF networks at the ensemble level. Fig. 3 shows that the probability to converge to a QFP in the approximated networks follows the same dependence on network size as the original SF ensembles. In particular, the significant difference between SFO and SFI, mapped to either an outgoing or incoming lumped hub, is captured by the approximation. These results are presented for a specific SFO ensemble, characterized by a fixed set of parameters (see figure caption for details).

**FIG. 3.**
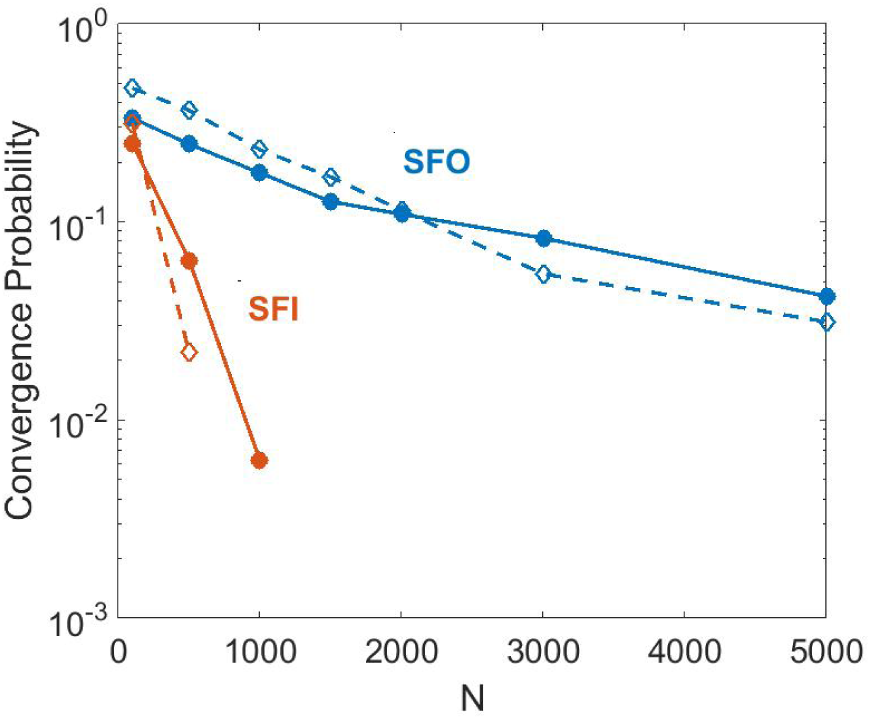
Dynamic properties of SFO/SFI lumped-hub approximation. Probability of convergence to a Quasi Fixed Point (QFP), for SFO and SFI (blue and red filled circles respectively; same data as in Fig. 1A). Their corresponding lumped-hub approximations are shown in blue and red open circles, with lumping parameter *m* = 4. Statistics and parameters as in Fig. 1.

We next ask whether our lumping approximation can reproduce the range of different behaviors across SFO ensembles with various power-law distributions. To answer this question we map a broad range of SFO ensembles to lumped-hub networks and compare their dynamic properties. Fig. 4 shows a scatter-plot of convergence probabilities for SFO networks vs. their lumped-hub approximations. Each point represents a specific SFO topology realization *T*, and the convergence probability is estimated as an average over realizations of connection strength *J*. It is seen that the convergence probability is largely retained in the mapping across a range of SF scaling parameter *γ* (Fig. 4A). In addition, Fig. 4B shows that the distribution of frozen core (FC) sizes, exhibiting a bi-modal shape dependent on ensemble parameters, is also very well captured by the lumping approximation. These results demonstrate that our lumped-hub approximation describes dynamic properties of the SFO network ensembles across a broad range of parameters.

**FIG. 4.**
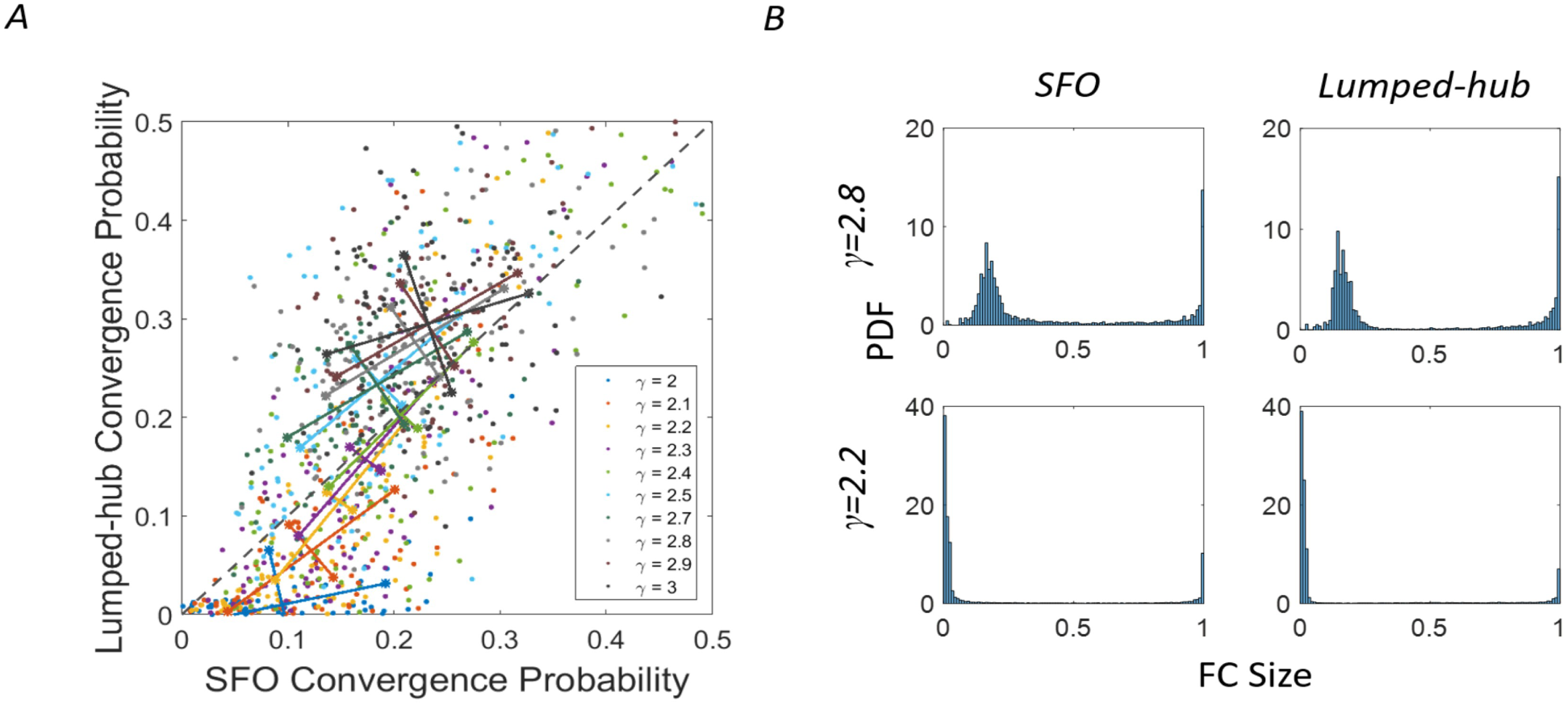
Lumped approximation captures SFO network properties for a range of power-laws.. **(A)** Probability of converging to QFP for SFO networks and their respective lumped approximations. For each SFO parameter *γ* (color coded), 100 topology realizations *T* were drawn, and the convergence probability was estimated from 50 realizations of the connection strength *J* and initial conditions *x*(0). Longer lines denote the direction of the linear fit to each *γ*, and the size of the perpendicular segment is the remaining variance after the fit. The line *y* = *x* is added for reference (dashed black). **(B)** Distributions of frozen core sizes for SFO networks and their lumped approximations for two power-law exponents. All SF distributions have *k*_*min*_ = 1 and the figure legend shows *γ* (see Methods for more details).

## IV. FEEDBACK ANALYSIS FOR THE LUMPED-HUB NETWORK ENSEMBLE

We have seen that despite its coarseness, the lumped hub approximation somehow captures an important feature of SF networks that strongly affects their dynamics. Can it be harnessed to gain a deeper understanding of these networks?

In particular we are interested in the significantly larger convergence probability in the SFO ensemble compared to the corresponding SFI. The essence of this difference becomes intuitively clear in the lumped-hub approximation. In lumped SFO networks, the hub coherently drives a macroscopic fraction of the bulk nodes. In contrast, in lumped SFI the hub receives a large number of inputs but only drives a small fraction (≈ *k*_*b*_) of the nodes. Previous work showed that intrinsically chaotic dynamics of homogeneous random networks can be suppressed by a coherent external drive [36]. Although here the hub is not external to the network, we argue that a similar suppression effect underlies convergence in SFO networks.

The lumped-hub network is described by two coupled equations of the bulk and the hub (Fig. 5):

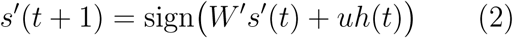

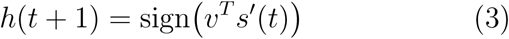

where *s*′ and *W*′ denote the bulk activity and connectivity respectively, the scalar *h* is the hub activity and the vectors *u, v* are the connections between the bulk and the hub.

**FIG. 5.**
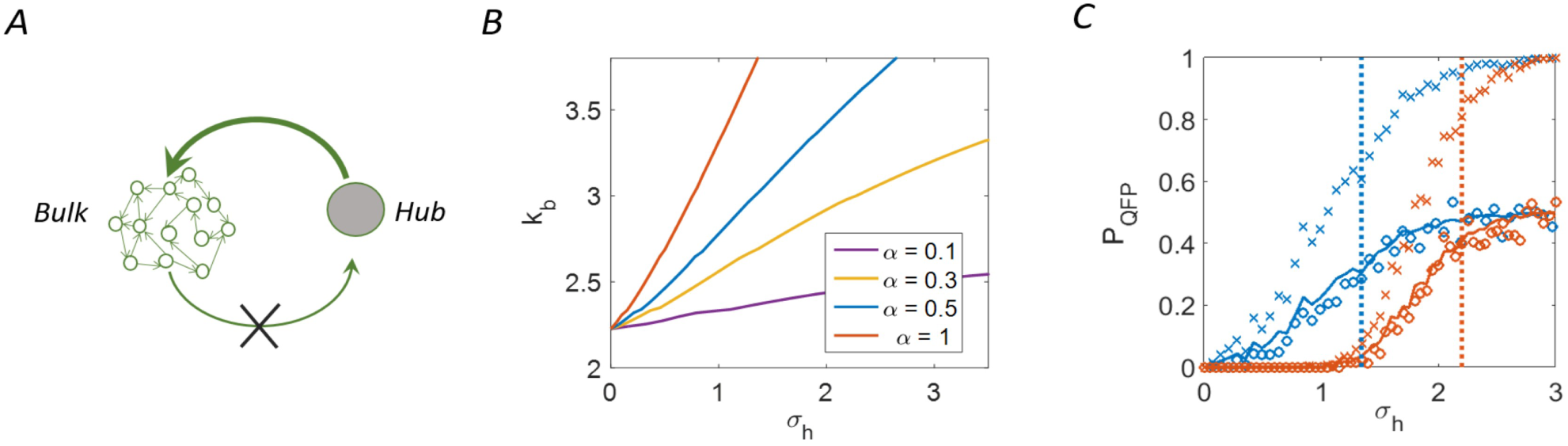
Feedback analysis for the lumped-hub network ensemble. (A) conceptual scheme of feedback analysis. The lumped-hub network contains a single hub (gray) connected to a homogeneous network representing the bulk (green). In the open loop setting, the hub is clamped to a fixed value while its incoming connections from the bulk are disabled (X). (B) Mean field theory predicts a threshold value of *σ*_*h*_ above which chaos in the bulk network is suppressed and the system is driven to a stable fixed point. The transition lines are plotted in the (*k*_*b*_, *σ*_*h*_) plane for various values of hub sparseness *α*. (C) The Probability of converging to a QFP for lumped-hub network ensembles computed as a function of hub strength *σ*_*h*_, for open-loop (top graphs, converging to 1) and closed-loop (bottom, converging to 1/2) settings. Two values of average bulk connectivity and sparseness are shown: *k*_*b*_ = 3, *α* = 0.5 (blue) and *k*_*b*_ = 5, *α* = 1 (red). Mean field theory predicts a phase transition at *σ*_*crit*_(*k*_*b*_, *α*), depicted by vertical lines of the corresponding color. The numerical results (symbols) confirm both the transition point and the factor of *half* due to closing the loop. *N* = 1500, and statistics is obtained from 239 realizations.

To facilitate the analysis, we consider a statistical ensemble characterized by three parameters: the lumped hub defines a sparseness *α*, the fraction of the network covered by its out-going connections; and a standard deviation *σ*_*h*_ of the strengths of these connections. The bulk in turn is characterized by the binomial parameter *k*_*b*_, derived from its average connectivity. In this simplification we lose the precise connectivity from the hub, which depended on the specific details of the original SF network. We shall see that several key properties are nevertheless captured by the simplified lumped hub ensemble with the three parameters (*α, σ*_*h*_, *k*_*b*_).

Consider first an auxililary setting – the “open loop” setting – where the hub is held at a fixed value. Effectively, connections from the bulk of the network to the hub are discarded, and it remains connected only through its outgoing links (Fig. 5A). The hub then acts as an input to the rest of the network, a simple situation that can be analyzed by mean field theory. Formally, we replace the dynamic variable *h* by a fixed value (+1 without loss of generality), namely Eq. (3) is substituted by *h*(*t* + 1) = 1. In this open loop setting, the regularizing effect of the fixed input competes with the recurrent dynamics of the bulk, and we ask under what conditions the system converges to a fixed point. Later, we connect the open-loop results back to a fully recurrent network including the hub as a dynamic variable [37].

To determine whether or not the system converges to a fixed point under input, we consider the time-evolution of the Hamming distance *d*(*t*) between two bulk trajectories 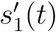 and 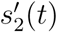. In the thermodynamic limit *N* → ∞, this evolution is deterministic and given by some function *f* (*d*) (see Ref. [18] and Appendix A here for derivations):

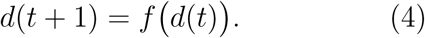

The external drive *h* can suppress chaotic activity and drive the network to a stable state *s*′* if and only if *d*\* = 0 is a stable fixed point of the mapping (4). This, in turn, requires the derivative of *d* at the origin, denoted by ∇*d*, to be smaller than *one*. We show in Appendix A that, within Mean Field Theory (MFT), the condition for this can be expressed as a threshold on the hub strength,

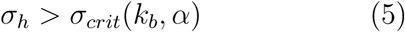

with *σ*_*crit*_ depending on the average bulk connectivity *k*_*b*_ and the sparseness *α*. Fig. 5B shows this critical line for two values of *α* in the (*σ*_*h*_, *k*_*b*_) plane. Importantly, the lines are distinct for different hub sparsity *α*, demonstrating that input strength cannot compensate for coverage; these are two separate properties of the hub that need to be accounted for correctly in the lumping approximation.

Fig. 5C illustrates the accuracy of our theory. Blue and red × symbols show simulated probability of convergence rising from zero to one for increasing *σ*_*h*_ for two value sets of (*k*_*b*_, *α*), with the critical values predicted by Equation (5) (vertical lines in corresponding colors). These results demonstrate both the qualitative prediction – a (smoothed) transition from zero to one in convergence probability, and the correct location of the transition at the predicted *σ*_*crit*_. We therefore conclude that MFT adequately models the open-loop convergence probability. Note that the critical driving strength increases indefinitely for large *k*_*b*_, in agreement with previous results [17, 18].

Returning now to the closed-loop network dynamics (2),(3), where feedback to the hub is restored and it is a dynamic variable with incoming and outgoing connections, we ask whether a stable attractor under external drive is consistently maintained as an attractor of the recurrent dynamics. Intuitively one may argue that, with probability 1*/*2 with respect to the random connections in the ensemble, the clamped value will be consistent with the hub’s input and the driven steady-state will also be an attractor of the dynamics. However, this intuitive argument only refers to existence of a closed-loop fixed point, and does not guarantee its stability. Therefore, beyond the open-loop stability criterion (5), the closed-loop setting requires to consider the recurrent dynamics of the hub. As long as the hub does not flip from its clamped value (*h* changes from +1 to −1), open and closed loop system dynamics coincide.

Consider a state *s*′, a small distance *d* = *δ* > 0 away from the fixed point *s*′*. For a stable fixed point we expect exponential convergence with a rate ∇*d*. Taking into account the finite size of the network, we estimate that convergence to a distance of less than one node (*d* < *N*^−1^) takes approximately

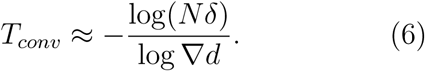

Preserving this stability in the full closed-loop network requires that the hub does not flip its sign during the time of convergence:

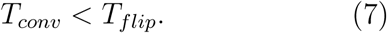

In Appendix D we show that the typical hub flipping time *T*_*flip*_ is approximately:

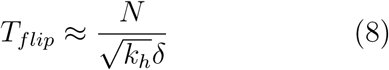

which implies that (7) holds for ∇*d* far enough from the phase transition. Furthermore, it is argued in the Appendix that even in the proximity of the critical value ∇*d* ≈ 1^−^ the corrections due to closing the loop are smaller than the error of the mean field approximation made in the open loop analysis. Taken together, these arguments show that all effects of closing the loop will reduce the probability of the system converging to a stable fixed point by half. Fig. 5C shows this is indeed the case. Circles depict closed-loop simulations of lumped-hub networks, and the matching lines show the smoothed result for open-loop divided by two.

## V. APPLICABILITY OF FEEDBACK ANALYSIS TO FINITE SFO NETWORKS

We now return to evaluate the accuracy of our mean field approximation on the original SF network ensembles. Making the connection between the theoretical results and empirical simulations entails two steps: first, for each SF network realization we define the lumped-hub parameters (*k, σ*_*h*_, *α*); second, these parameters are used to compute ∇*d* in MFT (Appendix A), predicting convergence for values smaller than one. Importantly, the first step is not a one-to-one transformation between the parameters of the SF ensemble and those of the lumped-hub ensemble: due to the large heterogeneity of SF networks, different realizations generally result in widely varying lumped-hub parameters and consequently in broadly distributed values of ∇*d*. An example of this distribution is shown for one SFO ensemble in Fig. 6A. In particular, lumped-hub parameters across the ensemble are such that ∇*d* is distributed on both sides of the phase transition (dashed vertical line, ∇*d* = 1).

**FIG. 6.**
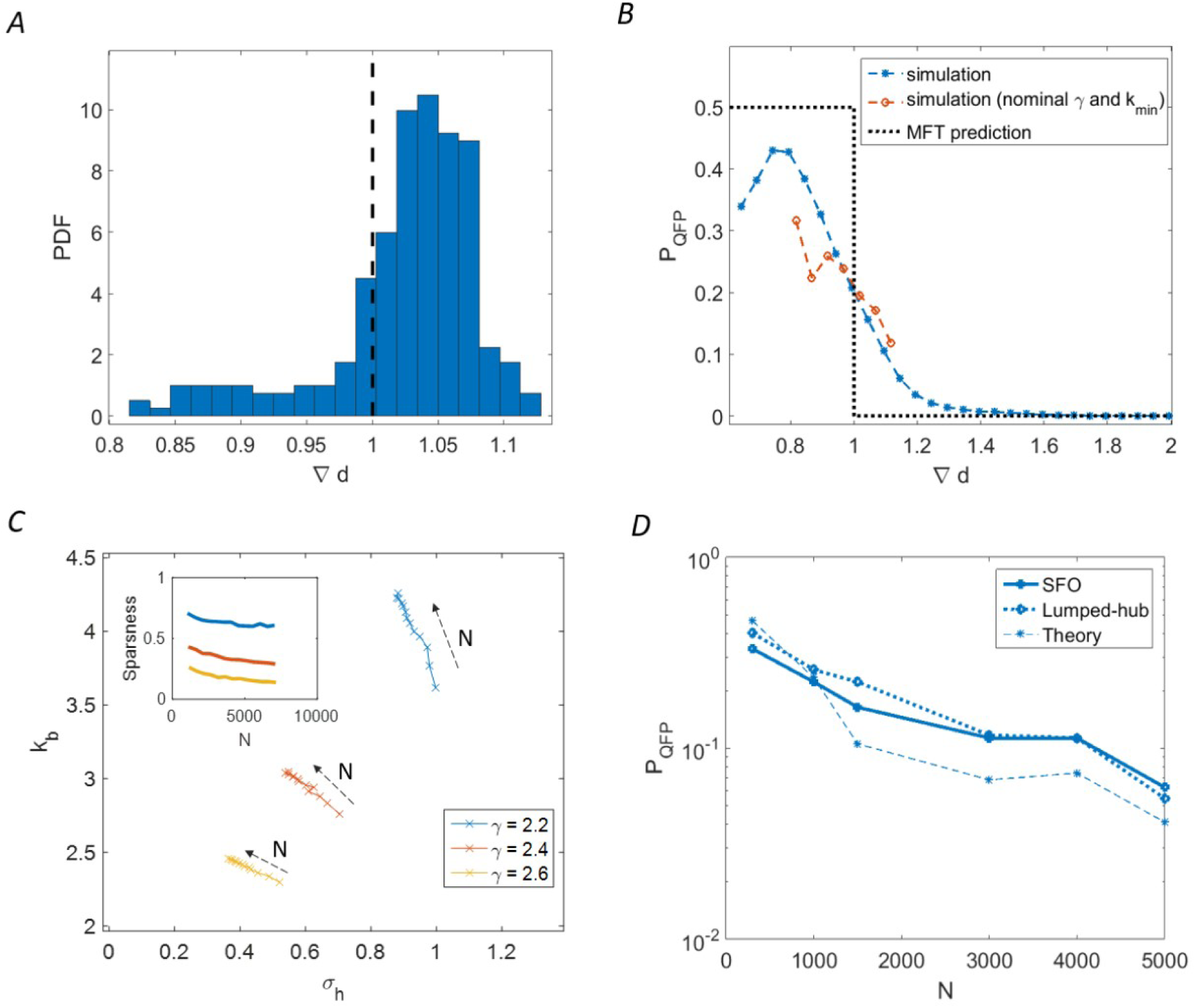
Comparing lumped-hub MFT prediction with SFO simulations. (A) Variability of lumped-hub parameters in a single SF network ensemble: histogram shows the distribution of ∇*d* as computed by MFT from the lumped-hub parameters, for a SF ensemble with *γ*_*out*_ = 2.4, *k*_*min*_ = 1. Location of MFT predicted phase transition, ∇*d* = 1, is depicted by a black line. (B) Transition in convergence probability predicted by MFT (black dashed line) and empirical calculation of *P*_*qfp*_ (colored symbols) with different *k*_*min*_ values in range 0.5 ≤ *k*_*min*_ ≤ 2 and *γ* = 2.2, 2.4, 2.6. For each set of parameters 256 network realizations of size *N* = 1500 were simulated. Simulation results were then binned by ∇*d* with bin width of 0.05. Bins with less than ten samples were omitted from the plot (restricting the standard error to 0.16 per point). A short red line shows the same data for the one ensemble with *γ* = 2.4, *k*_*min*_ = 1 only. (C) Effect of network size on average lumped hub parameters *k* and *σ*_*h*_. Increasing network size causes a gradual shift towards the unstable phase (see Fig. 5B). Sparseness remains approximately constant (inset). Colors show different *γ*_*out*_. (D) Effect of network size on convergence probability: comparing simulation, lumped-hub approximation and theory. Empirical data points represent 256 Monte-Carlo realizations for SF with *γ*_*out*_ = 2.4, *k*_*min*_ = 1.

Pooling together realizations of different power-law parameters opens a broader range of ∇*d* such that comparison to the theory is more meaningful. We simulated many SFO networks with a range of underlying power-law parameters and tested the prediction of a transition as a function of ∇*d*. Fig. 6B compares the theoretical prediction for the convergence probability (dotted black line) with the pooled simulation results (blue symbols); despite the pooling of a highly variable set of simulations, ∇*d* is revealed as a crucial determinant of convergence. The smooth sigmoid obtained follows the transition around the predicted value of ∇*d* = 1. For one SFO ensemble, the simulations follow the prediction but cover only a small range of ∇*d* (red symbols). A decrease in convergence probability is found for very low ∇*d* values, in contrast to the prediction; this effect will be discussed below.

Empirical convergence probabilities for SF networks also depend on network size (Fig. 1). This dependence can be partially explained by the lumped-hub approximation: for a given SF ensemble, we can follow the lumped-hub parameters averaged over the ensemble as a function of network size *N*. Fig. 6C shows that the average (*k*_*b*_, *σ*_*h*_) values progress towards the unstable regime as the network size increases, while the sparsity *α* is much less sensitive to network size; both these effects are consistent for several SF ensembles (colors). The implications of this trend can be seen in the dependence of convergence probability on network size, shown in Fig. 6D. Simulation results of SF networks and their corresponding lumped-hub approximations are shown together with the theoretical prediction, showing that most of the *N* -dependency is captured correctly.

## VI. LIMITS TO THE LUMPED HUB APPROXIMATION

The lumped hub approximation relies on the possibility of decomposing the network into a hub and bulk nodes. In turn, the phase transition predicted for lumped-hub networks relies on this decomposition and its relation to homogeneous networks driven by an external input. This approximation holds for a range of parameter values of practical interest. To understand the limits of validity for which the approximation breaks down, we consider several conditions that are required to hold.

First, the most highly connected nodes are being lumped together to one effective hub. The approximation thus neglects their internal dynamics, which should therefore be of little importance for convergence of the system. Ideally there should exist an interval of lumping numbers *m* across which ∇*d* remains largely unchanged. Fig. 7A shows that this is the case for SFO networks with *γ ≥* 2.4. For *γ* = 2.2 we see that ∇*d* depends more strongly on the choice of *m*, compromising the stability of the approximation. Indeed, Fig. 7B shows a large discrepancy between SF and lumped-hub behaviors for *γ* = 2.2.

**FIG. 7.**
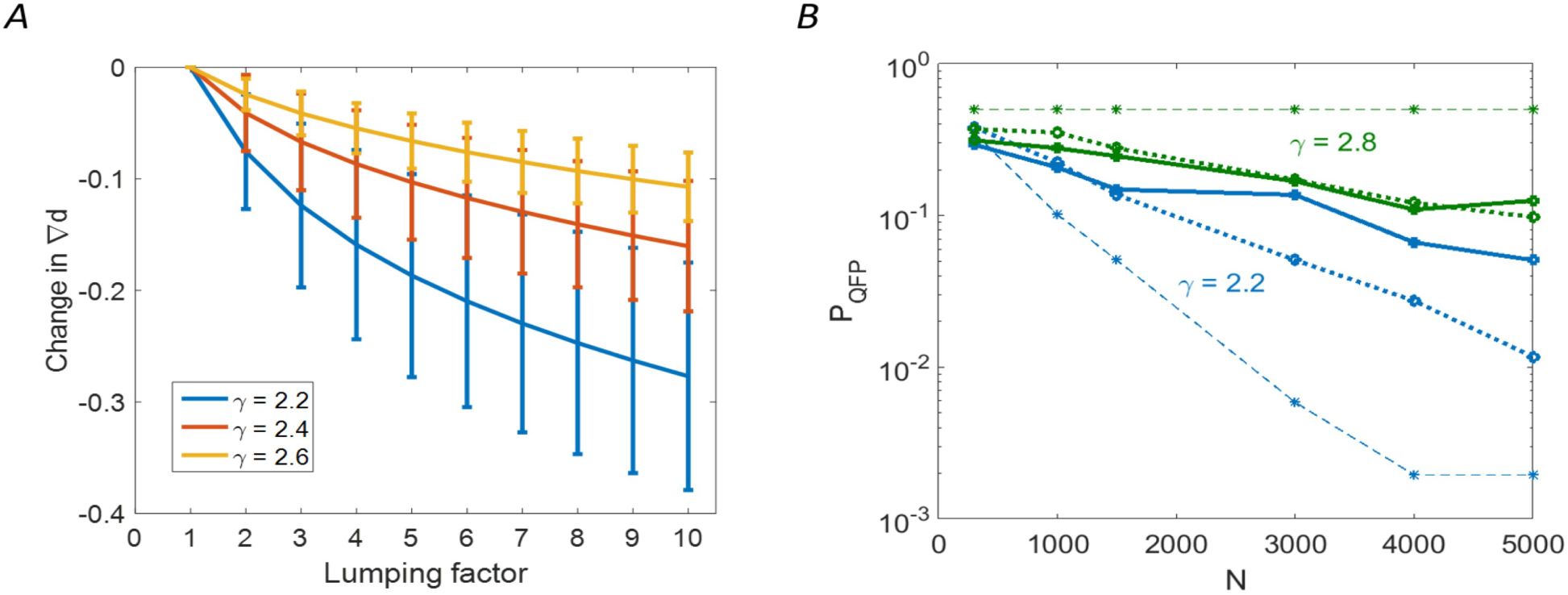
**A.** Sensitivity of lumped hub approximation to the number of leading nodes being lumped is shown for 1000 realisations of scale free topology with fixed parameters *k*_*min*_ = 1, *N* = 1500, and for three values of *γ*_*out*_ = 2.2, 2.4, 2.6. For each realization, the ∇*d* at *m* = 1 is taken as reference, from which the subsequent decrease in ∇*d* is measured **B.** Figure 6D is replicated for *γ*_*out*_ = 2.2, 2.8. In the large *γ*_*out*_ case, the lumped hub model holds, while mean field theory becomes invalid due to extreme sparseness. For low *gamma*_*o*_*ut* it is model that becomes inaccurate, due to sensitivity to choice of *m*.

Second, the network must remain *globally* connected. For very small mean connectivity, networks tend to split into a number of disconnected components. In this situation, the probability of an individual realization converging to a fixed point drops, while realizations with multiple fixed points emerge (Appendix C). We believe that this effect is responsible for the drop in convergence probability observed at extremely low ∇*d* (Fig. 6B).

In addition, for the approximation to be relevant, the effective external drive by the hub must be strong enough. As indicated in Fig. 5B, for a sparsely connected hub (*α* = 0.1) the transition between convergent and chaotic regimes is almost independent of hub strength. This can explain the failure of the theory for low *α* values, that arise for high *γ* values, as exemplified for *γ* = 2.8 in Fig. 7B.

Finally, by lumping *m* > 1 dominant hubs into a single node, we are limited to a single fixed point of the open-loop system (up to symmetry of the hub activation). If several combinations of the *m* hubs’ states can stabilize the open-loop system, the possibility of multiple fixed point pairs in a single realization emerges. According to [38] and Appendix C such a possibility should reduce the probability of convergence. This can partially explain the slight drop below the factor of 1*/*2 at Fig. 6B.

## VII. DISCUSSION

Broad connectivity profiles are a ubiquitous feature of networks. Understanding the dynamics arising from such topology, however, remains a great challenge. Guided by intuition from the physics of phase transitions, much attention has been given to the scaling of such distributions with network size (e.g. power-law and scale-free distributions). Real networks, however, are always of finite size – and our work highlights the benefits of focusing on this aspect. Our main hypothesis is that, for finite size networks, a small number of nodes has a disproportionate effect on the entire network. We suggest a simplified model − the lumped-hub approximation − that makes this hypothesis explicit by lumping these nodes into a single hub and replacing the rest of the network by a homogeneous bulk. This is a novel approach to finite size effects, different from the standard expansions around very small or very large networks. As such, it is suitable for intermediate network sizes, which are usually difficult to analyze. We showed that the lumped-hub approximation predicts the behavior of scale free networks at the ensemble level, explaining the change in convergence probability as a function of power-law exponent or network size. Furthermore, the approximation is also helpful at the single realization level, indicating that a large part of the variability between scale-free networks stems from the variability in their effective lumped-hub strength and coverage, and in the average bulk connectivity. We thus propose a three-parameter description of topology that is more predictive than the generative SF parameters.

The simplicity of the lumped-hub network ensemble, with a homogeneous bulk network and a single hub connected to all other nodes, allows analytic treatment of the problem. Specifically we applied feedback analysis, based on interpreting the hub as an external drive to the bulk, and then devising a mean field theory to determine convergence of the system. We then closed the loop while requiring consistency, allowing us to analytically estimate the probability for convergence. This analysis explains simply and intuitively why networks with outgoing hubs, and not those with incoming hubs, tend to converge to fixed points. We showed that, perhaps surprisingly, it is the stability of fixed points, rather than their abundance that underlies this effect.

The analytic treatment was developed for binary dynamic variables, but the empirical convergence properties and their dependence on outgoing hubs are shared also for a continuous variable model [27]. Therefore it is plausible that these properties reflect more generally the ability of outgoing hubs to promote convergence and to stabilize network dynamics.

We note the comparison of our results with previous work on Boolean networks with a randomly chosen Boolean function at each node [39], which found that the phase transition did not differ between SFO and SFI networks. By specifying a threshold function in our work the effect of a hub becomes coherent among all the nodes that it influences. In contrast, when randomizing over Boolean functions, the effect of the hub has no such coherence; it is this coherence which promotes convergence in recurrent dynamics with large hubs.

Our analysis was based on scale-free connectivity, but should apply for other topologies with broadly distributed out degree. In fact, our results indicate that the existence of a small number of hubs connected to a macroscopic fraction of the network nodes, rather than the precise shape of the distribution tail, is a crucial factor in shaping network dynamical properties. The existence of one such outgoing (but not incoming) hub was sufficient not only to induce increased convergence to stable attractors, but also to endow the ensemble with an extremely high heterogeneity among realizations.

From a biological viewpoint, our results illustrate how outgoing hubs − master regulators − can act as global coordinators of dynamics in gene regulatory networks. This role is expected to be realized under strong perturbations, in addition to the usual role of hubs as regulators of local specific circuits. Indeed, recent experimental work in bacteria suggests an organizing role for hubs in the emergence of gene expression patterns following strong rewiring perturbations [40, 41]. This idea, suggesting a dual role for the structure of gene regulation networks depending on conditions, awaits further experimental and theoretical investigation.

## METHODS

### Constructing Samples of Heterogeneous Network Ensembles

To construct network samples from an ensemble with a SFO dgree distribution, the adjacency matrix is constructed by first randomly sampling a sequence of *N* degrees from a scale-free distribution, and assigning a degree, *k*, from that sequence randomly to each node in the network. For each such node a set of *k* random outgoing connection is chosen. This procedure results in a scale-free outgoing distribution and Binomial incoming distribution as there are no constraint on the incoming distribution and the choice of incoming degrees is purely random. SFI ensembles where constructed by transposing SFO networks created as described above.

The scale-free sequences are obtained by a discretization to the nearest integer of the continuous Pareto distribution 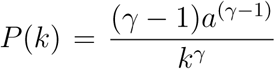. Binomial sequences are drawn from a Binomial distribution *P* (*k*) = ℬ (*N, p*), with 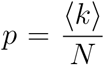, using MATLAB Binomial random number generator. Scale-free sequences are implemented by a discretization of the continuous MATLAB Generalized Pareto random number generators with Generalized Pareto parameters *k* = 1/(*γ* − 1), *σ* = *a/*(*γ* − 1) and *θ* = *a*.

### Probability of Convergence to fixed points and quasi fixed points

The networks’ frozen cores were determined by running the network dynamics for an initial convergence period of 4000 time steps, with 10% of networks nodes being updated at each step, and then measuring the fraction of nodes which were frozen within an additional time interval of 1000 steps (altogether 5000 time steps). Some simulations were run for double the time steps (a of total 10000 time steps, 2000 of which used for determining frozen core). No detectable difference was observed due to such longer times. Quasi fixed points were defined as states with a frozen core larger than 90%, while fixed points are networks with a frozen core of 100%. The probability of convergence to a fixed point or quasi fixed point was calculated by simulating the dynamics of a an ensemble of 500 networks, unless stated otherwise, and measuring the fraction of networks in the ensemble which reached the relevant frozen core criterion.

## VII. ACKNOWLEDGMENTS

This work was supported in part by the Israeli Science Foundation (grant number 346/16, OB; and grant number 155/18, NB)

## APPENDIX

## Appendix A: Suppression of chaos in the open-loop setting

We consider a lumped hub network ensemble where the effective hub is clamped at a fixed value and acts as an input to the rest of the nodes. Formally, we determine whether the input suppresses chaos by following the time evolution of the normalized Hamming distance between two trajectories *s*^1^(*t*) and *s*^2^(*t*), denoted by *d*(*t*). Suppose that *d*(*t*_0_) = 1*/N*, and assume, without loss of generality, that the states of interest *s*^1^, *s*^2^ differ at the first bit: 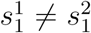 while 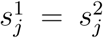 for 2 ≤ *j* ≤ *N*. At time *t*_0_ + 1 the distance *d* will be determined by the number of rows in *W* for which the first element changes its sign:

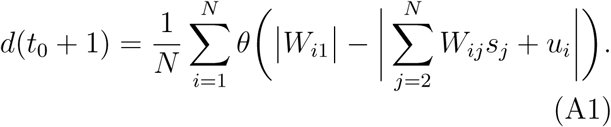

Equivalently, one may consider the derivative at the origin of the normalized distance *d*, given by:

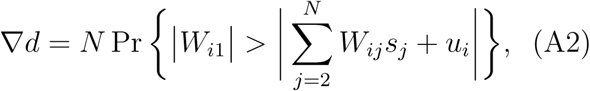

with the condition for stability being ∇*d* < 1. The inequality in curly brackets can be realized only if *W*_*i*1_ is nonzero; moreover the other matrix element can be grouped according to the number of nonzero elements in each row. This grouping results in the following convenient way to rewrite the last equation:

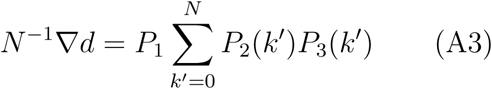

with terms *P*_1,2,3_ defined as:

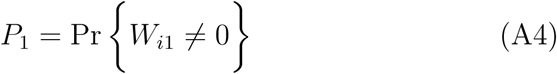

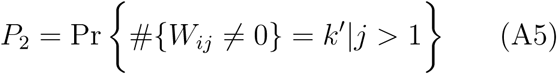

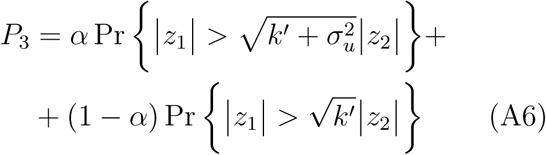

Here *z*_1_, *z*_2_ are i.i.d. normal Gaussian variables, as follows from the mean field approximation, and *i* is an arbitrary row. The terms *P*_1_ and *P*_2_ are determined by the network structure: *P*_1_ is simply the sparsity of the Binomial network which makes up the bulk nodes,

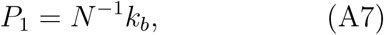

and the probability of a row in the network having exactly *k*′ nonzero entries is:

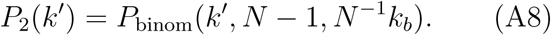

The only term which depends on the actual realization of weights is *P*_3_. To evaluate it we note that for a positive scalar *η*:

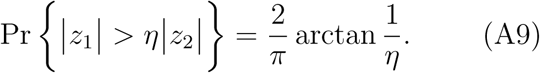

To derive this expression, one notes that in the |*z*_1_|, |*z*_2_|plane, we are interested in the portion of probability mass that lies below the line 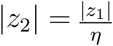. Since the probability density in this plane only depends on the distance from the origin, we simply compute the angle between this line and the |*z*_1_| axis and (A9) follows.

Combining all the terms together, and assuming large *N*, we arrive at:

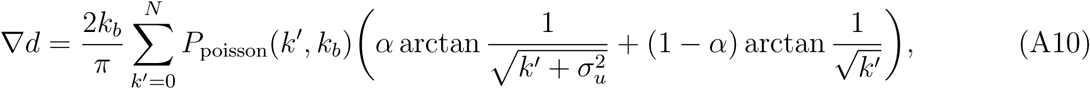

For *α* = 1 and large *k*_*b*_, the sum over *k*′ can be approximated by its expected value, and tanh by its argument leading to:

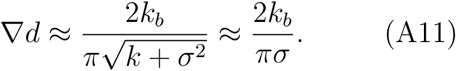

This expression accounts for the asymptotically linear relation between *k* and *σ* at the phase transition curve in Fig. 5B. Conversely, for sparse hubs with *α* < 1 there exists a *k*_*max*_ above which we have ∇*d* > 1 for any *σ*_*u*_.

### Appendix B: Quasi fixed points

Along with a possibility of converging to a stable fixed point another scenario exists where most network nodes are fixed but a small fraction is still toggling.

More specifically, a situation where all nodes except a small, interconnected clique are frozen

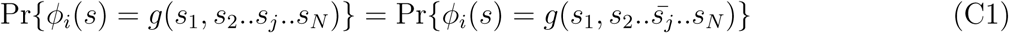

for any Boolean function *g*. In our case of threshold network, the condition (C1) is fulfilled

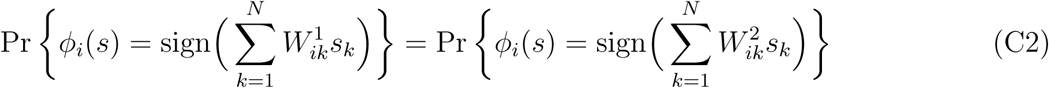

for Gaussian matrices *W*^1^ and *W*^2^ differing by inversion of a single element.

Unfortunately, the expectation, without any higher moments known, does not provide information about a typical case. Depending on the topology *T*, a typical realization of *W* might have *M* = *O*(1). Alternatively, there could be becomes typical. We refer to such a regime as a quasi fixed point (QFP). A threshold to define a QFP was set empirically at a value of 0.9 where a rise in probability of frozen cores starts (Fig. 1B).

### Appendix C: Fixed points are always there, regardless of topology

Defined by equation (1), our system turns out to be a special case of the Theorem in Ref. [38] which states that the expectation 𝔼*M* of number of fixed points *M* in random Boolean networks is *one*, subject to a condition on the distribution of random Boolean functions that holds in our setting. Specifically, to comply with [38] an ensemble of random Boolean functions *φ*_*i*_(*s*) defining the network dynamics via *s*_*i*_(*t*+1) = *φ*_*i*_(*s*(*t*)), must have a set of *neutral links*, removal of which renders the network acyclic. Here a link *j* → *i* between a pair of nodes is said to be neutral iff immediately because a few exceptional realizations with a huge number of fixed points while typically *M* = 0. For example, for *T* = *I* there are *M* = 0 fixed points with probability 1 − 2^−*N*^ and *M* = 2^*N*^ with probability 2^−*N*^. Conversely if *T* is a cyclic graph (e.g. cyclic permutation) then with probability of *half* network has two fixed points and it has *zero* fixed points in the complementary case. Although it is not immediately clear which scenario is relevant for SF networks, for lumped hub networks some insights are available.

For a lumped hub network with a strong hub, such that ∇*d* < 1 in the open loop setting, global convergence to a stable fixed point *s*′ = *s*\* is expected. If the consistency condition associated with relaxation of clamping of *h*(*t*) in (3) is met, then the closed loop system inherits the fixed point at *s* = *s*\*. Moreover, another fixed point emerges at *s* = − *s*\* corresponding to a drive of − 1 in the open loop system. Since the aforementioned condition is met with probability *half* we have an expected number of 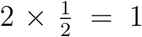 fixed points in agreement with [38].

In the opposite extreme case of *zero* drive from the hub, it follows from mean field theory that the network exhibits chaotic dynamics with any pair of state-space trajectories becoming asymptotically orthogonal. In particular, to comply with MFT, distinct fixed points should be either orthogonal to- or inverse of one another. We assume, without a rigorous proof, that orthogonality approximately implies independence. Namely, events of having a fixed point at *s* = *s*^1^ and at *s* = *s*^2^ are statistically independent for *s*^1^ ⊥ *s*^2^. Now, we recall that fixed points appear in pairs. Hence the number of *pairs* of fixed points is distributed binomially with parameters 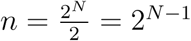 of possible pairs ±*s*\* of a state and its inverse, and a probability *p* = 2^−*N*^ that a particular pair of points is fixed. In the limit of *N* → ∞ this distribution converges to Poisson with parameter 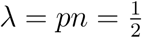:

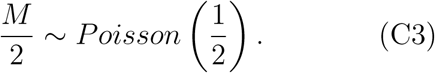

To validate this theory numerically, we performed an exhaustive search for fixed points in small, fully connected networks with 16 ≤ *N* ≤ 22 nodes. For *N* > 22 an exhaustive search was not feasible due to computational constraints. The results, shown in Fig. 8, depict an approximate match to the theory which is not perfect, with inaccuracies exceeding the standard error. We attribute these inaccuracies to the finite size of the networks, emphasizing that orthogonality of chaotic trajectories upon which our theory is built, is not achieved, even approximately, for values of *N* small enough for exhaustive search. Future work may include numerical experiments with larger and sparser networks using advanced search algorithms e.g. the method of [42].

**FIG. 8.**
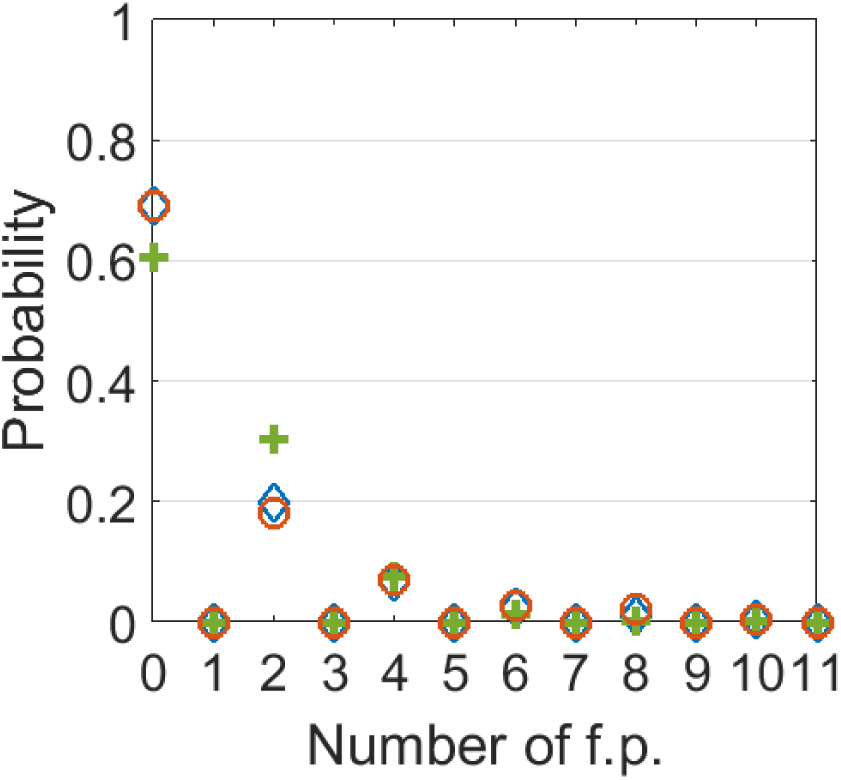
Probability of a given number of fixed points. ‘+’ – theoretical prediction. ∘ and – ◊ empirical estimates for networks of size *N* = 16 and 19 based on 10^4^ and 10^3^ Monte Carlo tries respectively.

For intermediate values of *σ*_*h*_ the picture is more complicated. Chaos is not suppressed but the symmetry between ±*s* is broken by the drive (in the open loop setting only, since once the loop is closed symmetry is restored). For *σ*_*h*_ ≪ 1 one might repeat our reasoning for *σ*_*h*_ = 0, this time with single fixed points rather than pairs thereof, and therefore with (C3) being replaced by:

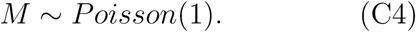

Numerical observations on small networks reported in table I tell us that while a sharp transition in the distribution of fixed points is observed once a drive is introduced, the distribution of *M* does not immediately match (C3). Here again we tend to attribute such a discrepancy with theory to finite size effects compromising the fixed point orthogonality.

**TABLE I.**
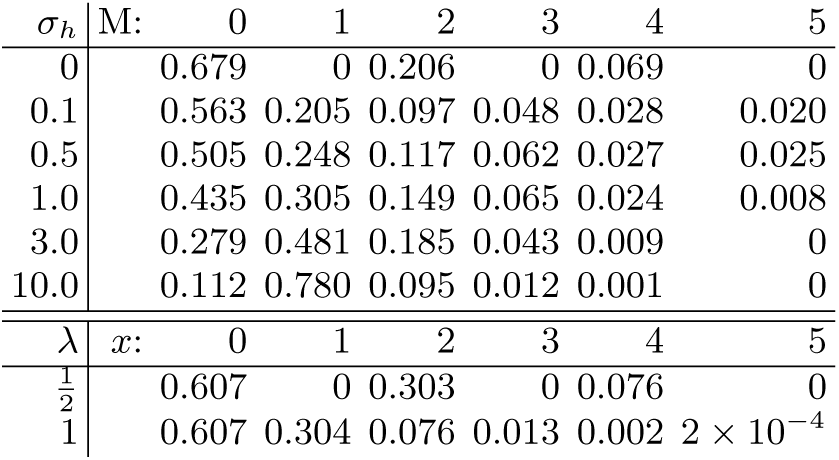
Numerical estimates for the distribution of *M* are shown as a function of external drive strength for ensemble of dense random networks of size *N* = 16 with elements gaussian i.i.d. Random gaussian drive is scaled by *σ*_*h*_. Poisson distribution with parameter 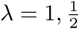 are provided for reference.

| $\sigma_h$ | M: | 0 | 1 | 2 | 3 | 4 | 5 |
| --- | --- | --- | --- | --- | --- | --- | --- |
| 0 |  | 0.679 | 0 | 0.206 | 0 | 0.069 | 0 |
| 0.1 |  | 0.563 | 0.205 | 0.097 | 0.048 | 0.028 | 0.020 |
| 0.5 |  | 0.505 | 0.248 | 0.117 | 0.062 | 0.027 | 0.025 |
| 1.0 |  | 0.435 | 0.305 | 0.149 | 0.065 | 0.024 | 0.008 |
| 3.0 |  | 0.279 | 0.481 | 0.185 | 0.043 | 0.009 | 0 |
| 10.0 |  | 0.112 | 0.780 | 0.095 | 0.012 | 0.001 | 0 |
| $\lambda$ | $x$: | 0 | 1 | 2 | 3 | 4 | 5 |
| $\frac{1}{2}$ | | 0.607 | 0 | 0.303 | 0 | 0.076 | 0 |
| 1 | | 0.607 | 0.304 | 0.076 | 0.013 | 0.002 | $2 \times 10^{-4}$ |

To conclude, we present a strong evidence, that the event of having a fixed point in the setting (1) has probability of order *one* and it is the fixed point stabilities rather than their existence that determines the convergence probabilities of dynamical numeric simulations like in Fig. 1.

Remarkably, this result is opposed to a case of continuous activation function (e.g. tanh) where an exponentially large amount of fixed points appear in the chaotic regime [43]. While an in-depth analysis of these settings in beyond the scope of the current work, we hypothesize that the difference between tanh and sign non-linearities follows from a fixed point at *zero*, which exists in the former case but is absent in the latter one.

### Appendix D: Estimation of hub flipping time in perturbed closed loop network

To obtain a, fairly coarse, lower bound on *T*_*flip*_ one may argue as follows: the hub has approxi-mately *k*_*h*_ = *m*⟨*k*⟩ = *O*(1) incoming connections whose weighted sum determines its sign. Let *H*_*in*_ denote the set of these nodes. Given a random perturbation of magnitude *δ* ≪ 1 to the fixed point *s*\* the probability for a single node in *H*_*in*_ to be flipped is

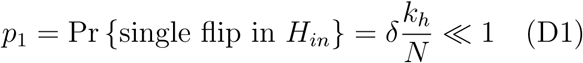

and the probability of *q* flips is 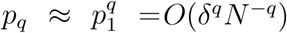 which can be neglected for small *δ*. Finally, by the arguments of Appendix A, the probability that the hub will flip due to a single node flip is 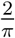 arctan 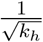 and consequently, the flip rate is given by:

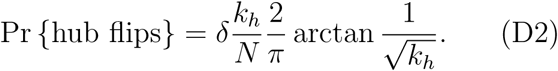

This corresponds to a typical flip time of:

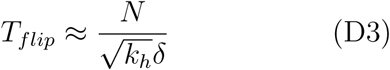

Hence, stability condition (7) clearly holds for − log ∇*d* of an order of *one*.

Even in the critical regime, where log ∇ → 0 is approached, condition (7) translates into

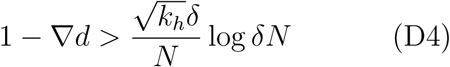

which for our typical setting of *N* = 1500, < *k*_*h*_ >≈ 20, and a reasonable small perturbed fraction of *δ* = 0.1 of the nodes translates into a requirement of:

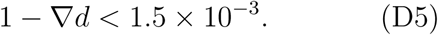

This is very close to criticality compared to other effects that may dominate inaccuracy of mean field calculations even in *open* loop setting. (Compare to Fig. 5 that depicts open loop and closed loop setting, and Fig. 6 which includes plot of convergence probability vs. ∇*d*).

